# Manipulation of alternative splicing of *IKZF1* elicits distinct gene regulatory responses in T cells

**DOI:** 10.1101/2024.11.05.622018

**Authors:** Lucia Pastor, Jeremy R.B. Newman, Colin M. Callahan, Rebecca R. Pickin, Mark A. Atkinson, Suna Onengut-Gumuscu, Patrick Concannon

## Abstract

Genome-wide studies have identified significant allelic associations between genetic variants in or near the *IKZF1* gene and multiple autoimmune disorders. *IKZF1*, encoding the transcription factor IKAROS, produces at least 10 distinct transcripts. To explore the impact of alternative splicing of *IKZF1* on the function of mature T cells, we generated a panel of human T-cell clones with truncating mutations in *IKZF1* exons 4, 6 or both. Differences in gene expression, chromatin accessibility, and protein abundance among clones were assessed by RNA-seq, ATAC-seq and immunoblotting. Clones with single targeting events clustered separately from double-targeted clones on multiple parameters, but overall, clone responses were highly heterogeneous. Perturbation of *IKZF1* splicing resulted in significant differences in expression and chromatin accessibility of other autoimmunity-associated genes and elicited compensatory expression changes in other IKAROS family members. Our results suggest that even modest alterations of *IKZF1* splicing can have significant effects on gene expression and function in mature T cells, potentially contributing to autoimmunity in susceptible individuals.

**Author Summary:** RNA sequencing has revealed an unexpectedly large population of alternative transcripts produced by most human genes, but the functional significance of such transcripts has been debated, with some authors arguing that they represent non-functional “noise” and others arguing that they are largely functional and greatly expand the potential human proteome. Here, we explore these issues for the transcription factor *IKZF1*, which produces numerous alternative transcripts in human T cells and is significantly associated with risk for type 1 diabetes and other autoimmune disorders. We introduce stop codons at multiple sites in alternatively spliced exons and evaluate transcript levels and chromatin accessibility in individually targeted clones, allowing us to assay the broad impact of alternative splicing of *IKZF1* in human T cells. Our studies reveal that perturbation of *IKZF1* splicing results in significant differences in gene expression, chromatin accessibility and protein production of other autoimmunity-associated genes and elicits compensatory expression changes in other IKAROS family members. Even modest alterations in *IKZF1* splicing had significant effects on gene expression and function in mature T cells, potentially contributing to autoimmunity in susceptible individuals and suggesting that transcript isoforms produced by alternative splicing can and do have functional impacts.

## Introduction

Studies of the human transcriptome by RNA-sequencing (RNA-seq) have revealed an unanticipated diversity of transcripts, suggesting extensive alternative splicing of both coding and non-coding RNAs. Estimates of the number of distinct human transcripts range as high as 300,000 (1). However, for the majority of these transcripts, their functional consequences, and for coding transcripts, their ability to produce meaningful levels of protein, remains largely unresolved and it has been argued that many alternative transcripts may be rare and/or of limited functional import (2, 3).

Transcription factors, which can regulate the expression of multiple genes are, arguably, an example of a class of genes for which alternative splicing could have broad, pleiotropic effects. Here, we explore the functional consequences of perturbing the distribution of protein isoforms that can potentially be generated by alternative splicing of the lymphoid transcription factor *IKZF1* in mature T cells. The *IKZF1* gene encodes IKAROS, the archetypal member of the IKAROS family of transcription factors that play critical roles in both lymphocyte development and function (4) (5, 6). *IKZF1* is encoded on chromosome 7 in a region that is associated with risk for several autoimmune diseases, including type 1 diabetes (T1D) (7–9) and systemic lupus erythematosus (SLE) (10), where it has been proposed as a candidate gene for these associations. In the case of T1D, *IKZF3* and *IKZF4*, are, like *IKZF1*, located in independent regions (on chromosomes 17 and 12, respectively) where there is also statistically significant evidence of association with disease risk (8, 9) suggesting a broader involvement of the IKAROS family of transcription factors in autoimmunity. IKAROS has been shown to regulate the expression of large numbers of genes in lymphocytes, including other genes implicated in risk for autoimmunity (11). Alleles at several genetic variants within the *IKZF1* gene are associated with variation in the expression of *IKZF1* as well as other genes located on multiple chromosomes, suggesting that these variants function as expression quantitative trait loci both in *cis* and *trans* (12). At least one of these variants, rs4917014, is significantly associated with risk of autoimmunity, specifically SLE (13).

Members of the IKAROS family of transcription factors share a conserved structure consisting of an N-terminal cluster of three or four zinc finger motifs forming a DNA binding domain and a second cluster of two zinc fingers at the C-terminal end of the protein that facilitate homo- and hetero-dimerization with other transcription factors, including other IKAROS family members (14, 15). Most members of the IKAROS family either produce, or have the potential to produce, multiple protein isoforms via alternative splicing. In the case of *IKZF1*/IKAROS, at least eight protein isoforms have been documented, and RNA transcripts with the potential to encode many additional isoforms have been reported (4, 16). These isoforms can be divided into two broad classes on structural and functional grounds: those that maintain both DNA binding and dimerization activity and those that lack the DNA binding domain while maintaining the dimerization domain (17, 18). These latter isoforms dominantly and negatively interfere with IKAROS function by competing for the binding of transcription factors. Mutations in *IKZF1* that favor the production of these isoforms are frequently observed in acute lymphoblastic leukemia (14, 18, 19).

Much attention has focused on the role of *IKZF1* in lymphoid development and on the effect of separating the DNA-binding and dimerization domains, as occurs in acute lymphoblastic leukemia. Here, we focus on the role of the zinc fingers that make up the DNA binding domain of IKAROS in mature T cells and the role of alternative splicing of *IKZF1* in modifying risk for autoimmunity. Specifically, we perturbed the distribution of *IKZF1* isoforms by creating frameshift mutations in exons containing zinc finger domains using CRISPR/Cas9 technology. Then, we explored the impact of these perturbations on the production of IKAROS protein isoforms and, more broadly, on chromatin accessibility and gene expression in cloned T cell lines by the Assay for Transposase-Accessible Chromatin (ATAC-seq) and RNA-seq analysis, respectively. Our results reveal surprising heterogeneity in the response of individual cells to manipulation of IKAROS isoform production, provide insight into the relative roles of individual zinc fingers within the DNA binding domain of IKAROS, and document the effects of altered splicing at *IKZF1* on the expression of downstream target genes, including those involved in autoimmunity.

## Results

### Targeting frameshift mutations to alternatively spliced *IKZF1* exons

We used CRISPR/Cas9 gene-editing technology to introduce frameshift mutations into exon 4 or 6 of *IKZF1* (reference transcript ENST00000331340.8) in Jurkat E6-1 cells using two distinct single guide RNAs (sgRNAs) for each exon. Cell clones were isolated by limiting dilution and the sequences of *IKZF1* exons 4 and 6 from individual clones were determined. Sixteen clones (of 30 with detected mutations) were selected for further analysis: seven (of 18) clones that had biallelic frameshift mutations in exon 4 (C5 – C11; **Fig 1, S1 Table**) and five (of 12) clones with biallelic frameshift mutations in exon 6 (C12 – C16). We also selected two clones with wild type sequences in both exons 4 and 6 (C1 and C2). Two additional Jurkat clones with wild type *IKZF1* sequences were selected from an independent targeting experiment (C3 and C4). One clone, C16, with a single nucleotide deletion in exon 6 on both alleles, was retargeted using the 4sg1 sgRNA for exon 4 (**S1 Table**). Eight clones containing biallelic frameshift mutations in both exons were selected from this double targeting experiment for further analysis (C17 – C24) (**Fig 1, S1 Table**).

**Fig 1.**
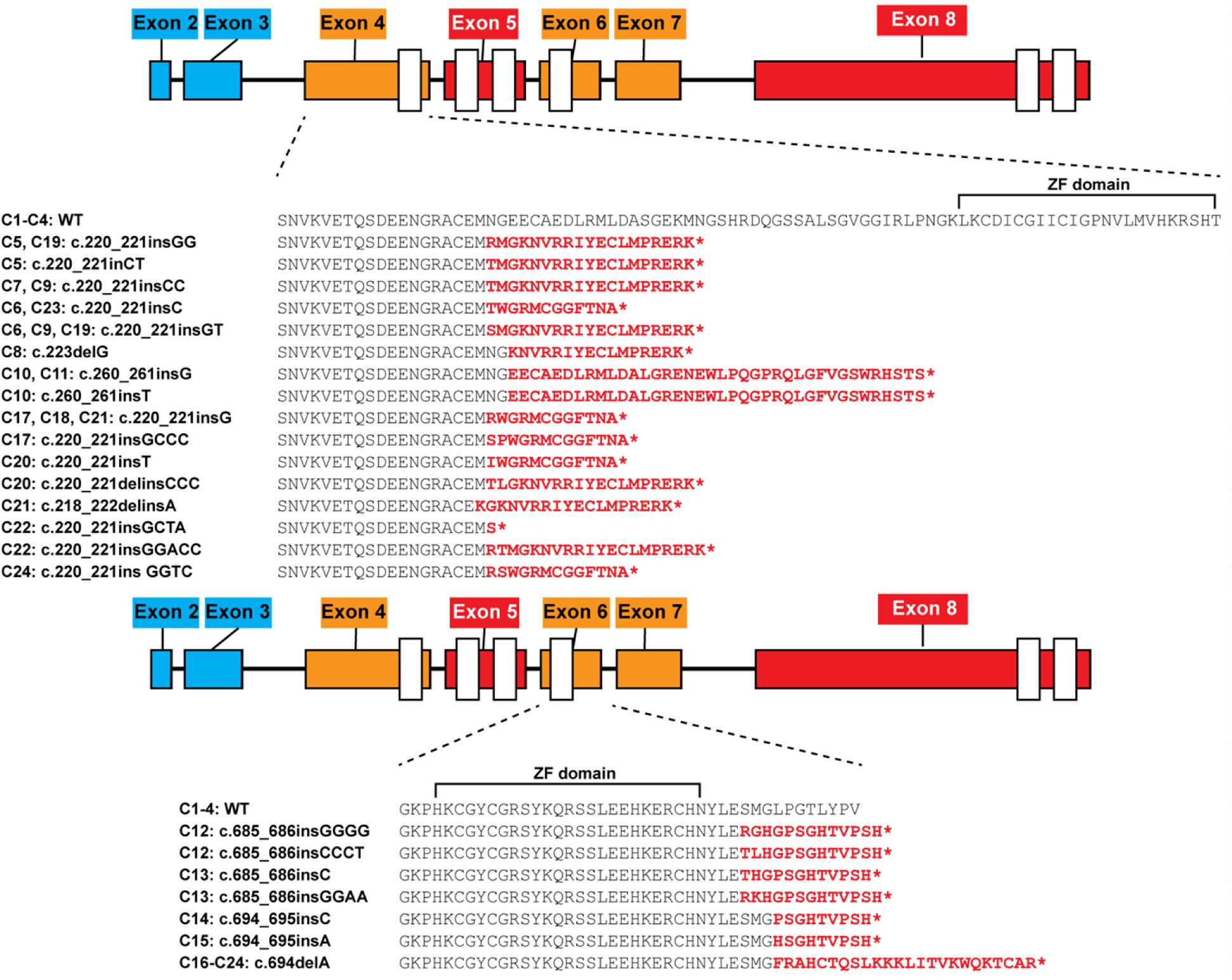
Predicted IKAROS proteins by clone. The consequences of the targeting events in Jurkat clones on the potential IKAROS isoforms produced are indicated. Amino acids depicted in red represent the predicted novel sequences resulting from frameshift mutations. “*” indicates a termination codon. The locations of the zinc finger (ZF) domains relative to the frameshift sequences are indicated by brackets labeled “ZF domain” above the amino acid sequences.

Exons are pictured as color boxes and numbered according to Ensembl transcript ENST00000331340.8. Exons alternatively spliced among the reported IKAROS functional isoforms are colored orange. Exons containing zinc fingers but that are not alternatively spliced are indicated in red. All other exons are colored blue. WT = wild type. Exons 4 and 6 of *IKZF1* contain 261 and 126 nucleotides, respectively, and the absence of either or both exons shortens but does not alter the translational reading frame of IKAROS. There are naturally occurring transcripts that lack these exons and produce alternative IKAROS protein isoforms (**S1 Fig**). The potential consequences of the frameshift mutations created here, with regard to the addition of amino acids encoded by the altered reading frame and the first termination codon that would be encountered are summarized in **Fig 1**. Additional details, including the sgRNAs used in the targeting of each clone and the mutations detected via DNA sequencing, are provided in **S1 Table**. The targeted sites in exon 4 precede the first zinc finger of the IKAROS DNA binding domain. Transcripts retaining this mutated exon 4 can only potentially produce a short amino-terminal protein lacking any DNA binding motifs. The frameshifts in exon 6 all occur immediately after the zinc finger domain. IKAROS protein translated from transcripts that retain the mutated exon 6 could contain an intact DNA binding domain but would lack the C-terminal half of the protein.

### Mutations in alternatively spliced *IKZF1* exons alter IKAROS isoform distribution

In order to assess the impact of exon targeting on IKAROS protein isoform expression, protein lysates were prepared from Jurkat clones and immunoblotted with antibodies directed against either the C-terminal (**Fig 2**) or N-terminal (**S2 Fig**) ends of IKAROS. Wild type clones, lacking mutations in *IKZF1*, expressed multiple IKAROS isoforms at consistent levels. In clones with frameshift mutations in exons 4 and/or 6, IKAROS protein isoforms containing sequences from these exons were not detected (**Fig 2A and 2B**), nor were novel N-terminal IKAROS isoforms that might result from premature termination due to the presence of frameshift mutations (**S2 Fig)**. Unlike wild type clones, exon-targeted clones displayed considerable heterogeneity in both the IKAROS isoforms (Iκ-1, Iκ-2, Iκ-3 and Iκ-4) produced and their levels of expression.

**Fig 2.**
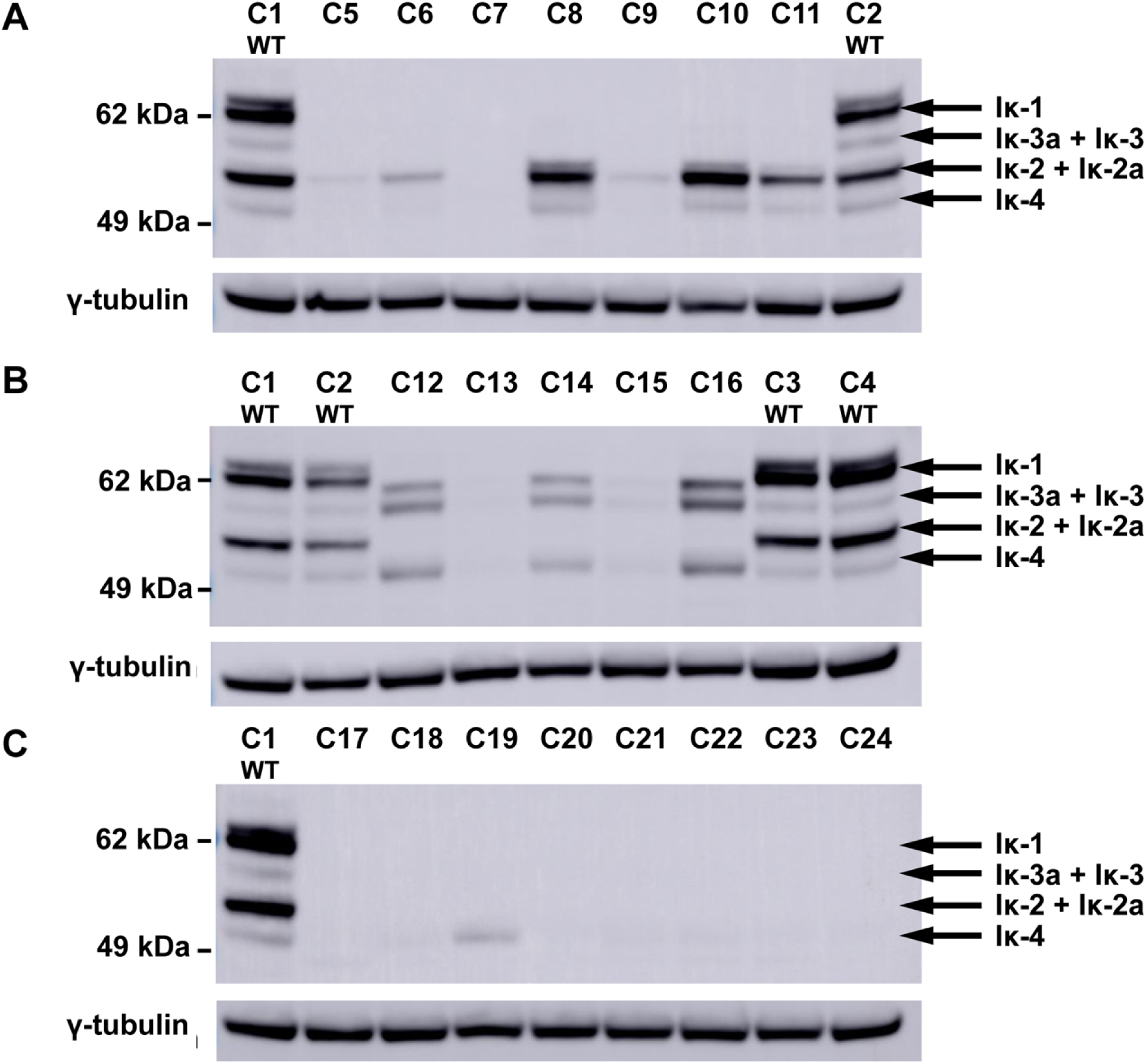
IKAROS protein expression in targeted clones. IKAROS protein expression as assayed by immunoblotting of clones with targeting events in exon 4 (A), exon 6 (B) and both exons (C), as compared to wild type clones (WT), using a primary antibody specific for the C-terminal end of IKAROS. The positions corresponding to migration patterns of IKAROS splice isoforms, Iκ-1, Iκ-2/Iκ-2a, Iκ-3/Iκ-3a and Iκ4 are indicated on the right. Results obtained with an N-terminal targeted primary antibody are provided in **S2 Fig**. The blots were subsequently stripped and probed with an antibody specific for γ-tubulin as protein loading control. Black arrows indicate the expected sizes of IKAROS isoforms containing the N-terminal zinc fingers encoded by exon 5 (i.e. Ik-1, Ik-2, Ik-2a, Ik-3, Ik-3a, Ik-4; **S1 Fig**).

Expression levels did not correlate with the guide RNA used for targeting, nor with the nature or location of the resulting frameshift mutations. Three exon 4 targeted clones (C5, C7 and C9) and two exon 6 targeted clones (C13 and C15) displayed little or no detectable IKAROS expression even though protein isoforms generated by the skipping of exons 4 and/or 6 were observed in both wild type clones and other exon-targeted clones. Two exon 4 targeted clones, C8 and C10, expressed the Iκ-2 isoform, which lacks exon 4, at greater than wild-type levels. Similarly, three exon 6 targeted clones (C12, C14 and C16) expressed levels of the IKAROS isoforms Iκ-3, Iκ-3a and Iκ-4 at increased levels relative to wild-type clones. Clones with mutations in both exons 4 and 6 produced little or no detectable IKAROS, with the exception of clone C19, which produced approximately wild-type levels of an isoform corresponding to either Iκ-4 or Iκ-4a (**Fig 2C)**.

### Transcriptomic analysis reveals differences in *IKZF1* exon usage after gene targeting

RNA extracted from all *IKZF1*-targeted clones at the same time that protein lysates were prepared was subjected to RNA-seq. Overall, *IKZF1* transcript levels did not differ significantly between wild-type and exon-targeted Jurkat clones (P > 0.05). However, when clones were grouped by target exon, a gradient of *IKZF1* expression (wild type > exon 4 targeted > exon 6 targeted > exons 4 and 6 targeted) was observed with the exon double targeted clones having significantly lower expression of *IKZF1* than wild type clones or exon 4 targeted clones (P = 0.018 and P = 0.017, respectively). The usage of individual exons exhibited the same gradient as overall *IKZF1* gene expression except for two infrequently used and alternatively spliced exons located between exons 3 and 4 (**Fig 3A**). These two exons are unique to *IKZF1* transcript isoforms that lack sequences encoding the DNA binding domain. These transcripts were enriched in double targeted clones (ENST00000440768, blue transcript) and reduced in wild type clones (ENST00000492782, purple transcript, **Fig 3B**). Variation in the abundance of some transcripts was consistent with the targeted exon. For example, the expression of transcripts lacking exon 4 (ENST00000698574 and ENST00000439701, orange and teal transcripts respectively) were elevated in exon-4 targeted clones, particularly those with higher levels of overall IKAROS protein expression (C8, C10 and C11) and lower in exon-6 targeted clones as compared to wild type clones.

**Fig 3.**
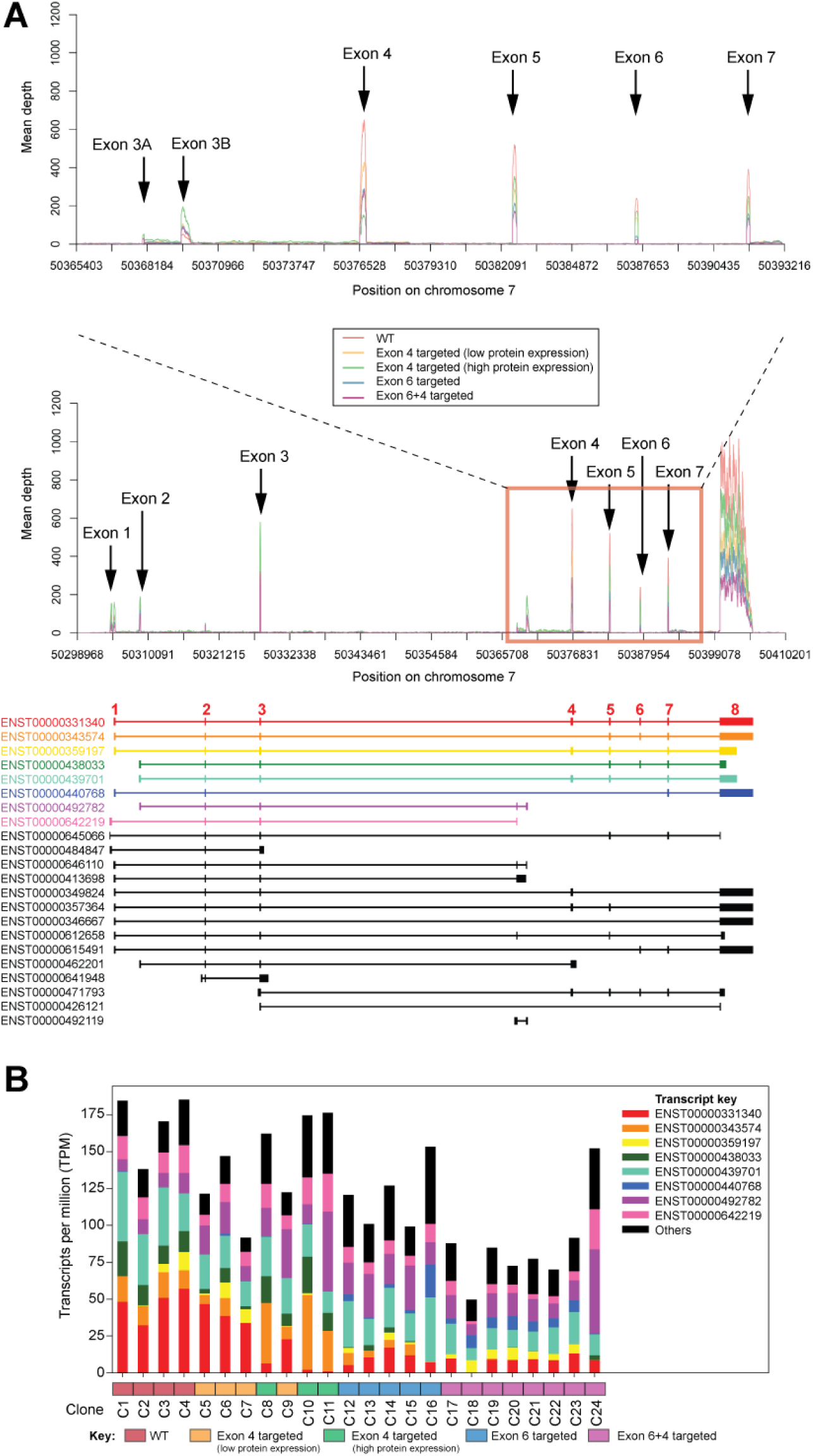
*IKZF1* exon usage and transcript expression in targeted clones. **A**) Transcriptomic analysis of the *IKZF1* exon usage by exon-targeted groups. Wiggle plots depict the mean coverage of the *IKZF1* gene relative to physical location in the studied clones grouped as wild type (red line); exon 4 targeted clones with either low (orange line) or high protein expression by immunoblotting (green line); exon 6 targeted clones (blue line) and double targeted exon 6+4 clones (purple line). The full *IKZF1* locus (± 1kb) and the chromosomal region containing exons 4 through 7 are shown with *IKZF1* transcript models. Numbered exons are annotated in red according to transcript ENST00000331340.8. The x axis indicates the position on chromosome 7 in nucleotides from version GRCh38.p13 of the human genome assembly. **B**) Expression levels of *IKZF1* transcripts by clone. The different transcripts are color coded by transcript identifier as in panel A and their expression in TPM is represented in the stacked bars for each clone. Total bar height corresponds to the total *IKZF1* gene expression for the indicated clone. Colors in the horizontal axes represent the different clone groups according to the exon(s) targeted and IKAROS protein expression.

We considered the possibility that the heterogeneity in IKAROS protein expression observed in the targeted Jurkat clones might result, in part, from the creation and use of new splice donor or acceptor sequences by the gene targeting events or the activation of cryptic splice sites. A limited number of RNA-seq reads consistent with the usage of novel splice sites were detected. However, the usage of novel splice sites was, at best, a very minor contributor to heterogeneity in *IKZF1* expression.

### Perturbation of *IKZF1* splicing affects global gene expression and chromatin accessibility

Using the same set of *IKZF1*-targeted Jurkat clones, we jointly analyzed gene expression data generated by RNA-seq with genome-wide profiling of chromatin accessibility generated using the ATAC-seq to assess the broader functional impact of perturbing *IKZF1* splicing beyond the immediate impact on IKAROS expression. Gene-targeted clones were analyzed as (1) the combined set of all 20 clones, (2) grouped by specific target exon(s) and level of IKAROS protein expression, or (3) by the number of targeted exons without regard to their specific identities (WT, one targeted exon, two targeted exons). This last condition is equivalent to grouping by the number of zinc finger motifs present in the DNA binding domain of IKAROS (e.g., 4, 3 or 2, respectively). A total of 20,875 expressed genes and 227,594 ATAC-seq peaks between IKZF1-targeted clones and wild type clones were compared. We identified 1,886 differentially expressed (DE) genes and 3,287 differentially accessible peaks (DAPs) comparing wild type clones and one or more of the 3 groupings of *IKZF1*-targeted clones (FDR-corrected *P* < 0.05). Based on these statistically significant features, the clones clustered into groups that largely reflected the number, rather than the specificity, of their gene-targeting events and hence the number, but the not the identity, of the zinc finger motifs present (**Fig 4**). Clones with a single targeting event in either exon 4 or 6 were interspersed and clustered adjacent to the wild-type clones. Despite the evidence from immunoblotting that some singly-targeted clones expressed little or no detectable IKAROS protein, these clones did not cluster together in the expression analysis and were interspersed among the other exon 4 and exon 6 single-targeted clones (**Fig 4A**). Double-targeted clones clustered together with the exon 6 singly-targeted parental clone from which they were derived (C16), differing from other singly targeted clones in both gene expression (**Fig 4A**) and chromatin accessibility (**Fig 4B**).

**Fig 4.**
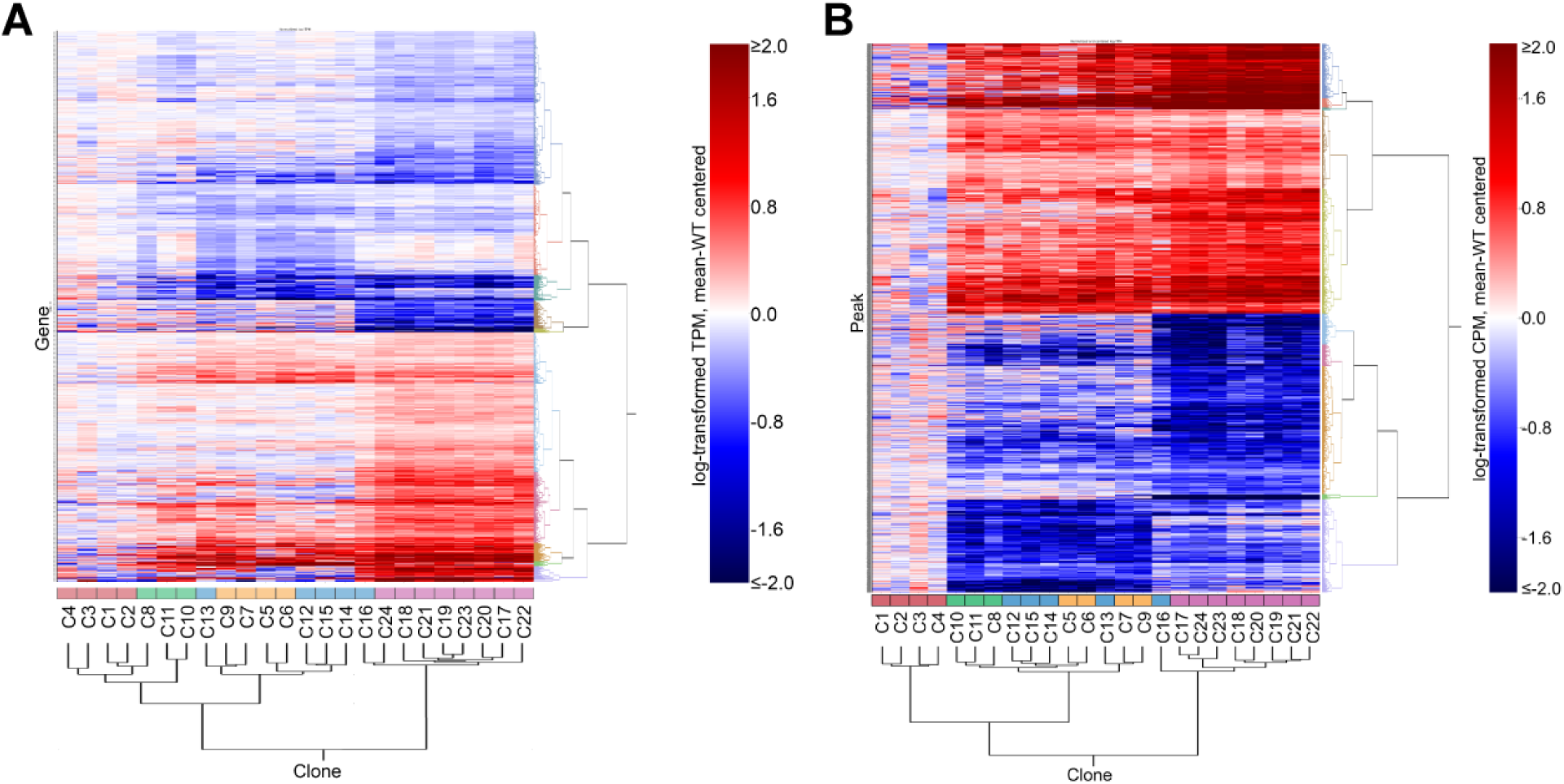
Differential gene expression and chromatin accessibility between wild type and *IKZF1*-targeted clones. (A) Heatmap showing the transcript levels of significant differentially expressed genes among the cloned Jurkat lines. (B) Heatmap showing peak levels in significant differentially accessible genes among the cloned Jurkat lines. Red boxes = wild-type clones, green boxes = exon 4-targeted clones with high protein expression, yellow boxes = exon 4-targeted clones with low protein expression, blue boxes = exon 6-targeted clones, purple boxes = exons 6 and 4-targeted clones.

Given that all single exon-targeted clones displayed largely the same profiled response (**Fig 4**), we hypothesized that it is the number of zinc finger motifs in the DNA-binding domain of IKAROS that is the major determinant of transcriptional and epigenetic changes induced by gene-editing of *IKZF1*; i.e. IKAROS function is dependent on the capability of producing functional isoforms with either four (WT clones: C1-C4), three (exon 4 or 6 single-targeted clones: C5-C16), or two (exon 4 and 6 double-targeted clones: C17-C24) zinc fingers in the core DNA-binding domain and that the specific sequences from either exon 4 or exon 6 have a negligible impact on IKAROS function compared to the overall difference in the number of zinc fingers. To formally test this hypothesis, we compared the gene expression and chromatin accessibility of: (1) high-versus low-expressing exon 4-targeted clones; (2) the combined set of exon 4-targeted clones versus exon 6-targeted clones; (3) the differences in number of targeted exons (WT vs. single-targeted, WT vs. double-targeted, single-targeted vs. double-targeted).

We categorized ATAC-seq peaks as genic if they were proximal (≤ 2kb) to the body of at least one gene (91,615 of 227,594 detected peaks; 40.3%) or intergenic if they were located greater than 2 kb from the nearest transcriptional start site (TSS) or transcriptional termination site (TTS) (135,979 of 227,594 detected peaks; 59.7%). We further defined differentially accessible (DA) genes as any gene having one or more proximal differentially accessible peaks (i.e., located within the gene body or within 2kb of the start or termination sites for transcription). We then examined the expression of genes with proximal ATAC-seq peaks among *IKZF1*-targeted and WT clones to identify genes that were significantly impacted by perturbation of *IKZF1* splicing. When comparing the gene expression levels between exon 4 targeted clones that displayed either low or high levels of IKAROS expression by immunoblotting, there were 138 DE genes (FDR-corrected *P* < 0.05), of which 12 genes were expressed consistently in the exon 4 clones with high levels of IKAROS (high-expressing exon 4 mean TPM > 1, low-expressing exon 4 mean TPM < 1, FDR-corrected *P* < 0.05; **Fig 5A**). However, none of these genes had differentially accessible peaks between high- and low-expressing clones, nor was any gene differentially accessible between these clones (**Supplemental Note**). No differences in either chromatin accessibility or gene expression were observed between exon 4-targeted clones and exon 6-targeted clones (**Fig 5A**). Comparison of WT and single-targeted clones yielded a few DE genes (83 DE genes; **Fig 5A**). However, when comparing double-targeted clones with WT or single-targeted clones, 1,212 and 2,887 DE genes were identified, respectively (**Fig 5A**).

**Fig 5.**
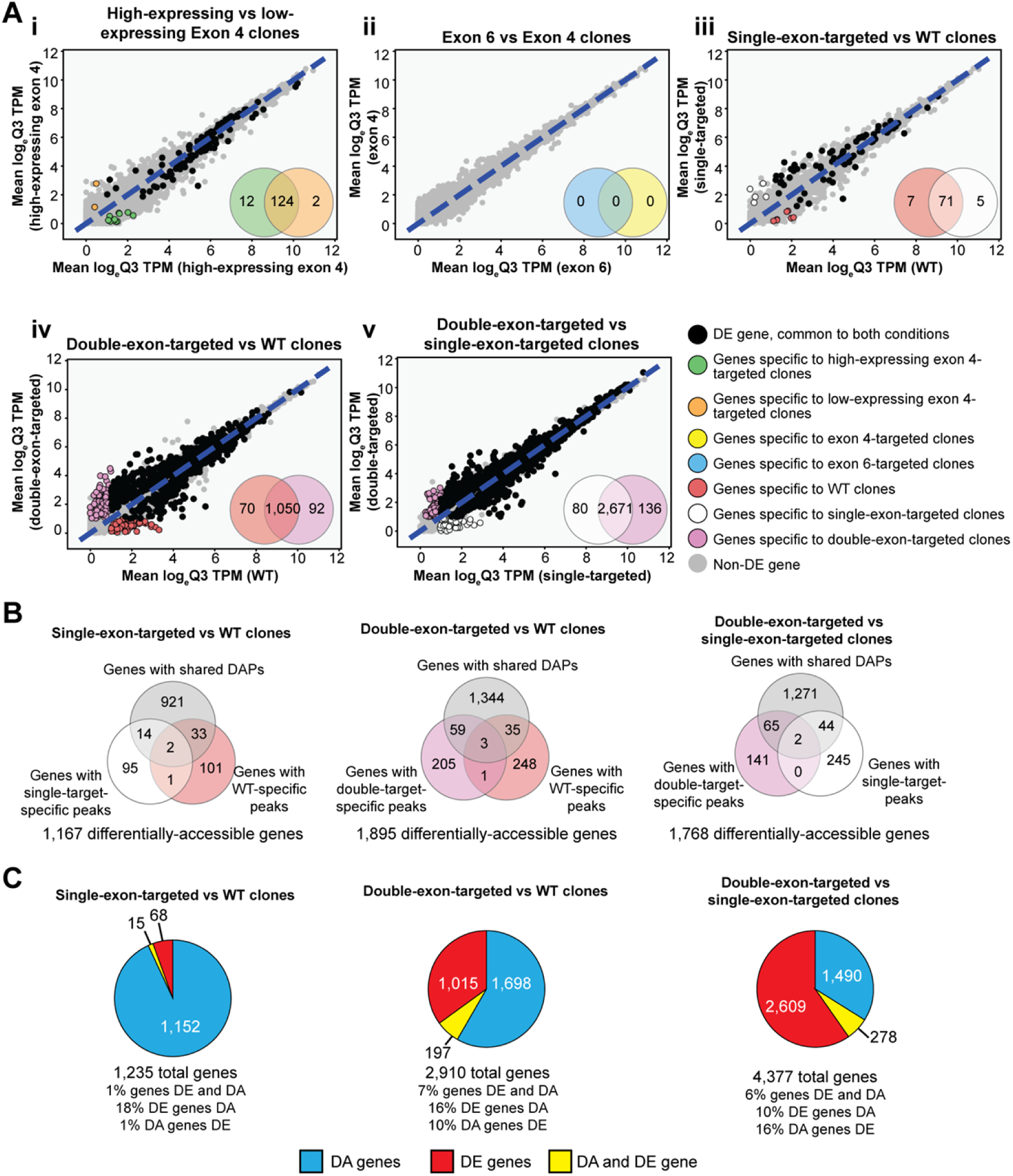
Transcriptional and chromatin accessibility analysis of *IKZF1*-targeted clones. (**A**) comparison of gene expression between (i) high protein-expressing exon 4-targeted clones and low protein-expressing exon 4-targeted clones, (ii) exon 6-targeted clones and exon 4-targeted clones, (iii) WT clones and single exon-targeted clones, (iv) WT clones and double exon-targeted clones, and (v) single exon-targeted clones and double exon-targeted clones. Venn diagrams report gene detection (mean logTPM ≥ 1) in each comparate and the genes detected in both comparates. The log-transformed TPM for each condition is plotted on the x- and y-axis. Black points represent significant differentially expressed (DE) genes detected in both conditions. Genes with significantly different expressions detected in only one condition are represented by colored points corresponding to their specific comparate. (**B**) Classification of genes with comparate-specific peaks and common, differentially accessible (DA) peaks between WT, single-targeted clones, and double-targeted clones. (**C**) Distribution of genes by differential features (DA genes, DE genes, DA and DE genes) between WT, single-targeted clones, and double-targeted clones.

Differences in chromatin accessibility between WT, single-targeted, and double-targeted clones were more frequent than observed for gene expression. In contrast to the relative paucity of DE genes, there were 1,167 DA genes between WT and single-targeted clones, 1,895 DA genes among WT and double-targeted clones, and 1,768 DA genes between single-targeted and double-targeted clones (**Fig 5B**). The majority of DA genes (76-83%) had at least one quantitatively different DAP (FDR *P* < 0.05) detected in both clone types compared (WT vs. single, WT vs. double, single vs. double) (**Fig 5B**), indicating that most differences in DA genes arise from differential accessibility at the same position/locus/region/site rather than from the presence or absence of distinct peaks.

We then assessed the overlap between DA and DE genes in *IKZF1*-targeted and WT clones. The number of genes, both DA and DE, in each comparison, was modest (1-7% of genes), and few DE genes were also considered DA (10-18% of DE genes; **Fig 5C**). Specifically, 15 of 83 DE genes were DA between WT and single-target clones, 197 of 1,212 DE genes were DA between WT and double-target clones, and 278 of 2,887 DE genes were DA between single-targeted and double-target clones (**Fig 5C**). The remaining genes were either DA or DE, for which the distribution varied between comparisons (e.g., the proportion representing DA genes) varied from 28% (double-vs. single-targeted clones) to 93% (WT vs. single-targeted clones).

We assessed the overlap in response (DA or DE) between the three clone group comparisons (WT vs. single-targeted, WT vs. double-targeted, and single vs. double-targeted) (**Fig 6**). Significant genes common to all three comparisons comprised only 3% of DA genes (108 of 3,484 genes; **Fig 6A**) but increased to 36% (1,238 of 3,484 DA genes) when considering genes DA in at least two comparisons. Less than 1% (8 of 3,359) of DE genes were common to all three comparisons, increasing to 24% (815 of 3,359 DE genes) when considering genes DE in at least two comparisons, with most of these shared DE genes arising from comparisons involving double-targeted clones (**Fig 6B**). No obvious patterns emerged when all 6,210 genes significantly affected by the targeting of *IKZF1* were categorized on the basis of differential accessibility and/or differential expression for each of the three comparisons; the most frequent responses tended to be comparison- and/or assay-specific (**Fig 6E**; **Supplemental Fig 3**) and relatively few genes were both DE and DA (N = 427 genes). Only a single gene – *FRMD4A* – was significantly different for both assays in all three comparisons. However, when considering any gene with a statistically significant difference in expression or chromatin accessibility, we note that many of the genes responsive to *IKZF1* targeting are in common between each of the comparisons (**Fig 6D**).

**Fig 6.**
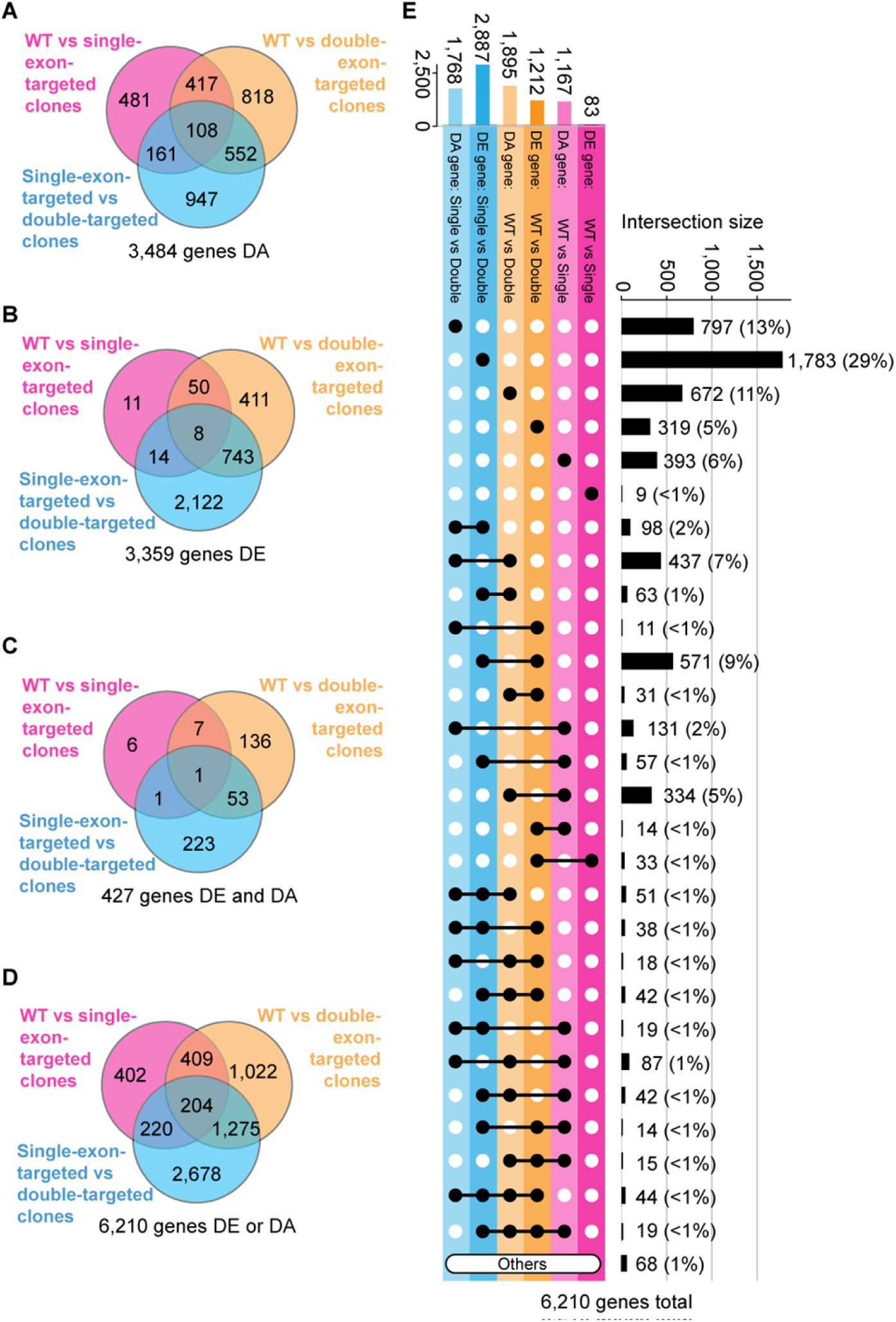
Comparison of differentially-accessible (DA) and/or differentially-expressed (DE) genes between WT, single-exon-targeted clones, and double-exon-targeted clones: (A) DA genes only, (B) DE genes only, (C) genes classed as both DA and DE (D) genes classes as either DA or DE. (E) Upset plot comparing statistically significant genes between WT, single-targeted, and double-targeted clones for each assay (chromatin accessibility, gene expression). Intersections with less than 10 members (excluding genes only DE between WT and single-exon-targeted clones) are grouped into the “Others” category. See **S4 Fig** for an extended version of this figure.

We also considered differential expression and differential accessibility as equal responses to editing events in *IKZF1* and compared the sets of affected genes for each of the three comparisons (**Fig 6D**). Few affected genes were common to all three comparisons made (204 of 6,210 (3%) total affected genes) and most affected genes were specific to one comparison (402 (magenta area only) + 1,022 (yellow area only) + 2,678 (blue area only) = 4,102 of 6,210 (66%) genes comparison-specific; **Fig 6D**). Genes affected by only a single *IKZF1* targeting event were a modest fraction of the total genes responding to the targeting of *IKZF1* (402 of 6,210 (6%) total affected genes, **Fig 6D** (magenta area only)) whereas clones with targeting events in both exons 4 and 6 elicited the greatest response (**Fig 6D**). About half of the genes affected by targeting a single *IKZF1* exon were distinct from those genes affected by targeting both exons 4 and 6. As the largest difference in terms of affected genes was observed between clones with a single targeted exon and clones with two targeting events (4,377 of 6,210 (70%) of the total set of affected genes; **Fig 6D**), this suggests that the response to the loss of a single zinc finger is distinct from that of the loss of two zinc fingers in the core DNA binding domain of IKAROS.

### Perturbation of IKAROS isoform distribution results in a broad response affecting multiple immune associated genes

We considered the possibility that perturbing the distribution of IKAROS isoforms might elicit compensatory changes in the expression or chromatin accessibility of other members of the IKAROS family. *IKZF4* and *IKZF5* were excluded from our analyses as their expression was undetectable or invariant, respectively, across the *IKZF1*-targeted clones (**Supplemental Note**). Among the remaining family members, *IKZF2* was both significantly DE and DA between WT clones and double-targeted clones (**Fig 7A**). *IKZF2* has three DAPs clustered within intron 4 of its main isoform (ENST00000434687.6) whose average peak heights followed a linear relationship with the number of targeted exons in *IKZF1* (**Fig 7A, S5A Fig**). Average *IKZF2* transcript levels followed a similar pattern, resulting in significant differences in expression between single-targeted and double-targeted clones (**Fig 7A**). Immunoblotting revealed that double-targeted clones express more Helios, the transcription factor encoded by *IKZF2*, than single-targeted clones; expression of Helios in WT clones was more variable (**Fig 7A**). *IKZF3*, which encodes the protein Aiolos, displayed a similar pattern to *IKZF2* in terms of differential accessibility and differential gene expression, (**S5B Fig**). The core DNA-binding domains of Helios and Aiolos share homology with the DNA-binding domain of IKAROS and given that all three proteins recognize an overlapping set of DNA motifs (15, 20), the observed changes in expression and chromatin accessibility of *IKZF2* and *IKZF3* may reflect compensatory responses to reductions in IKAROS function or changes in isoform distribution resulting from the targeting of *IKZF1* exons 4 and/or 6.

**Fig 7.**
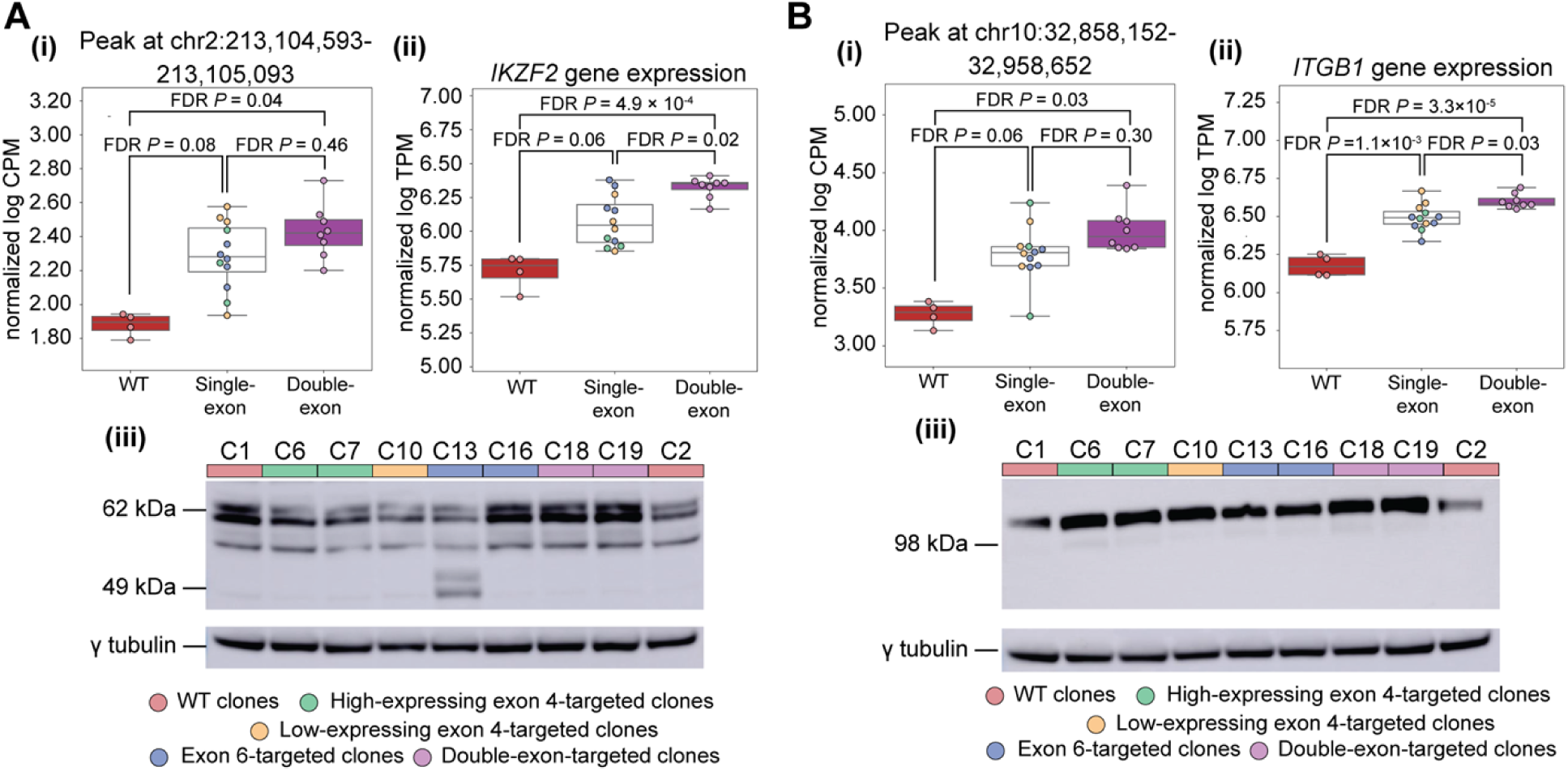
Effects of IKZF1-targeting on downstream regulated genes. The effects on two genes, *IKZF2* (A) and *ITGB1* (B), are shown in terms of chromatin accessibility **(i)**, gene expression **(ii)**, and protein levels **(iii)**. A representative example of changes in chromatin accessibility for each gene is presented here. Additional regions significantly differentially accessible (FDR *P*<0.05) and other example genes are presented in the extended **S5 Fig**. Red boxplots = WT clones; white boxplots = single-targeted clones; purple boxplots = double-targeted clones; pink/red dots = WT clones, green dots = low protein-expressing exon 4-targeted clones; yellow dots = high protein-expressing exon 4-targeted clones; blue dots = exon 6-targeted clones; purple dots = double-targeted clones.

Given the genetic associations observed between alleles of genetic variants located in or near IKAROS family members and risk for autoimmune disorders such as T1D and SLE (8, 13), we tested if genes located within established risk loci for autoimmune disorders were more likely to be among the genes that were DE or DA in our study. We found that SLE risk genes were enriched among DE genes when comparing double-targeted and single-targeted clones than were non-SLE genes (93 of 563 (17%) SLE risk genes; 2,794 of 32,825 (9%) non-SLE genes; χ^2^ = 44.92, DF = 1, *P* = 2.1 × 10^-11^), and also, to a lesser extent, in comparison of double-targeted clones and WT clones (39 of 563 (7%) SLE risk genes; 1,173 of 32,825 (4%) non-SLE genes; χ^2^ = 17.80, DF = 1, *P* = 2.5 × 10^-5^). Similarly, T1D risk genes were more frequently DE between double-targeted and single-targeted clones compared to non-risk genes (31 of 209 (15%) T1D risk genes; 2,856 of 33,179 (9%) non-T1D genes; χ^2^ = 10.19, DF = 1, *P* = 1.4 × 10^-3^). When considering all genes associated with autoimmune disease risk, we also observed that more autoimmune risk genes were DE compared to non-risk genes between double-targeted and WT clones (167 of 2,475 (7%) autoimmune risk genes; 1,045 of 30,913 (3%) other expressed genes; χ^2^ = 74.26, DF = 1, *P* = 6.8 × 10^-18^), and also between double-targeted clones and single-targeted clones (365 of 2,475 (15%) autoimmune risk genes; 2,522 of 30,913 (8%) other expressed genes; χ^2^ = 125.95, DF = 1, *P* = 3.2 × 10^-29^). SLE, T1D, and other autoimmune risk genes were not overrepresented among genes with a proximal DAP for any one specific comparison. However, when DAP-proximal genes from all three comparisons (WT vs. single-targeted, WT vs. double-targeted, single-targeted vs. double-targeted), we observed that T1D risk genes (13 of 209 (6%) T1D risk genes, 753 of 33,179 (2%) other genes; χ^2^ = 14.46, DF = 1, *P* = 1.4 × 10^-4^), SLE risk genes (32 of 563 (6%) SLE risk genes, 734 of 32,825 (2%) other genes; χ^2^ = 29.35, DF = 1, *P* = 6.0 × 10^-8^), and autoimmune risk genes in general (127 of 2,475 (5%) autoimmune risk genes, 639 of 30,913 (2%) other genes; χ^2^ = 95.99, DF = 1, *P* = 1.2 × 10^-22^) more frequently had a nearby DAP. Overall, considering all genes that were either DE or had a proximal DAP from any of the three comparisons, we found that T1D risk genes (47 of 209 (22%) T1D risk genes, 3,906 of 33,179 (12%) other genes; χ^2^ = 22.85, DF = 1, *P* = 1.8 × 10^-6^), SLE risk genes (134 of 563 (24%) SLE risk genes, 3,819 of 32,825 (12%) other genes; χ^2^ = 78.50, DF = 1, *P* = 8.0 × 10^-19^), and autoimmune risk genes in general (515 of 2,475 (21%) autoimmune risk genes, 3,438 of 30,913 (11%) other genes; χ^2^ = 205.99, DF = 1, *P* = 1.0 × 10^-46^) were more likely to be DE or DA.

Among the 427 genes that were both significantly DE and DA (FDR *P*<0.05) for at least one comparison of I*KZF1*-targeted clones (**Fig 6C**) is *ITGB1,* which encodes the beta-1 subunit of integrin, making up part of a membrane receptor in lymphocytes that regulates a variety of cellular processes involving cell adhesion and recognition (21). Four DAPs are located close to *ITGB1*, and as observed for *IKZF2*, there is a linear relationship between peak height at these DAPs and number of targeted exons in *IKZF1*-targeted clones (**Fig 7B**; **S5C Fig**). One of these peaks, located at chr10:32958054-32958554 also overlaps a reported IKAROS ChIP-seq peak (**S5C Fig**), suggesting that *ITGB1* expression may be directly regulated by IKAROS binding. It has been previously reported that the expression of the mouse homolog of *ITGB1* (*Itgb1*) is increased when exon 5 of mouse *Ikzf1* is deleted (22). In human Jurkat cells, targeting of exons 4 and/or 6 of *IKZF1* yielded a similar effect, with the average expression of *ITGB1* increasing with the number of exons targeted (**Fig 7B**). *ITGB1* was significantly DE in all three comparisons of *IKZF1*-targeted clone groups (WT vs single-exon, WT vs double-exon, single-exon vs double-exon; FDR *P*<0.05) (**Fig 7B**). ITGB1 protein expression was also elevated among targeted clones compared to WT (**Fig 7B**).

Responses to *IKZF1* targeting like that of *ITGB1* were also observed for other immune-related genes such as *IL15*, where average peak heights of three DAPs (one DAP overlapping the transcriptional start site of *IL15* and two DAPs overlapping the first exonic region) significantly increased with number of targeted *IKZF1* exons, as did overall gene expression (**S5D Fig**). Statistically significant differences in expression or accessibility of *IL15* were observed when WT or single-targeted clones were compared to double-exon-targeted clones (**S5D Fig**).

## Discussion

There is increasing evidence that genetic risk variants for various autoimmune disorders act through effects on transcript abundance, resulting in differences in overall protein expression levels or the distribution or nature of protein isoforms that a gene can produce (12, 23, 24). However, for most human genes, the functional significance of the production of multiple transcripts remains unresolved and its biological significance is a subject of controversy. In the current study, we focused on the role of alternative splicing, in mature human T cells, of the transcription factor IKAROS, encoded by *IKZF1*.

The members of the IKAROS family of transcription factors have a relatively conserved primary structure and are known to produce multiple transcripts via alternative splicing that result in the production of distinct protein isoforms. Much of the interest in IKAROS has focused on dominant negative isoforms that retain the C-terminal dimerization domain while lacking DNA binding activity and whose over-expression is associated with leukemia (14, 18, 19). To probe the consequences of alternative splicing of *IKZF1* on IKAROS DNA binding, we introduced truncating mutations into exons 4 and 6 of *IKZF1*, each of which is alternatively spliced and encodes zinc finger motifs that make up part of the DNA binding domain of IKAROS. We then generated and characterized multiple cloned cell lines containing different truncating mutations and combinations of mutations on the same genetic background. We observed significant heterogeneity among these cloned cell lines in both the distribution of IKAROS isoforms expressed and their relative levels of expression. As splice sites were not specifically targeted, clones should be capable of producing the normal distribution of *IKZF1* transcripts, although transcripts that included exons containing frameshift mutations might be underrepresented due to nonsense-mediated decay. Therefore, the IKAROS protein isoforms that could be produced by any specific clone would be limited by which exons contained frameshift mutations. However, even in clones where the *IKZF1* targeting resulted in the same or comparable frameshift mutations, there were significant differences in both *IKZF1* transcript expression and IKAROS protein isoform distribution. Clones with frameshift mutations in exon 4 were approximately evenly divided between those that expressed levels of IKAROS comparable to wild type, albeit with an altered distribution of isoforms, and those that produced little or no IKAROS protein detectable by immunoblotting with multiple antibodies. These clones differed largely in the ratio of the canonical full-length *IKZF1* transcript that includes exon 4 (ENST00000331340) and a second transcript (ENST00000343574), lacking exon 4, produced. Surprisingly, there were also multiple examples of targeted clones in which the abundance of specific IKAROS isoforms was increased relative to the same isoforms in wild type clones (e.g., isoforms Iκ3 and Iκ4 in clones C12, C14 and C16 or isoform Iκ2 in clones C8 and C10).

While only the two central zinc finger motifs, encoded in exon 5 of *IKZF1* are considered essential for DNA binding activity and nuclear localization, the elimination of protein isoforms that include either of the remaining flanking zinc finger motifs encoded in exons 4 and 6 might be expected to alter the DNA binding specificity or affinity of IKAROS in cells containing mutations in these exons (4, 15). We considered multiple models to explain the impact of the mutations in *IKZF1* on overall gene expression and chromatin accessibility, including grouping of clones by the specific exons targeted. However, we found that the clones clustered primarily based on the number of zinc finger motifs potentially available (4, 3 or 2) independent of the identities of the specific zinc fingers. This result differs from prior observations in mouse models in which the homologous exons were individually deleted. In these mouse models, distinct but overlapping sets of biological functions and Ikaros binding sites, assessed by ChIP-seq, were observed (4).

However, the current study and that by Schjerven et al. differ in several ways that may explain the disparate conclusions regarding the specificity of zinc finger function in IKAROS. In the mouse studies, T cells develop and mature in an environment influenced by the Ikzf1 mutations while the current study employs a human T cell line, albeit transformed, that mimics many of the functions of normal human T cells (25). The approaches also differ, employing deletions that result in the absence of the targeted exons as opposed to frameshift mutations that truncate protein isoforms containing those exons. The heterogeneity, in terms of overall IKAROS abundance, protein isoform distribution and transcript distribution, observed among Jurkat clones that differ only modestly in the nature of the mutations introduced in the targeted exons, suggests either a stochastic response to mutagenesis of these exons or a surprising sensitivity to the specific mutations induced.

While the association of *IKZF1* with leukemia reflects its role in lymphocyte development, the associations with autoimmune disorders, which have received less attention, likely reflect the role of IKAROS in the function of mature T and B cells. Common genetic variants associated with altered expression of *IKZF1* have been associated with risk for SLE (10, 13) while similar variants located near *IKZF1*, *IKZF3* and *IKZF4*, each of which is encoded on a different chromosome, are independently associated with risk for a second autoimmune disorder, T1D (8, 9). Rare deleterious coding variants in *IKZF1* have also been associated with various autoimmune, typically B cell mediated, phenotypes (26, 27). At the same time, alternative splicing, which, as demonstrated here and in prior studies, is a key characteristic IKAROS family members, has been found to mediate risk for autoimmunity at specific disease-associated genes such as *PTPN22*, *UBASH3A*, *SIRPG*, *IFIH1* and *CTLA4* (23, 24, 28–32). Globally, the set of all genes residing within chromosomal regions with significant genetic associations with one or more autoimmune disorders is enriched for alternative splicing (23). Finally, dysregulation of the RNA binding proteins involved in mediating splicing has been reported in both T regulatory cells and pancreatic Beta cells from subjects with T1D (33, 34). The current study adds to the accumulating evidence implicating alternative splicing in mediating genetic risk for autoimmunity. The set of genes that were either differentially expressed or differentially accessible upon manipulation of IKAROS protein isoform distribution via CRISPR/Cas9 targeting was significantly enriched for genes located in regions associated with autoimmunity, overall, and specifically those associated with either SLE and T1D. Both SLE and T1D are specifically associated with common genetic variants proximal to the *IKZF1* gene. The effects on T1D risk might be further augmented in patients by the significant genetic associations present at *IKZF2*, whose expression and accessibility is modified by IKAROS isoform usage as demonstrated here (**Fig 7A** and **Supplemental Fig 5A**). Thus, in this work, we not only documented the effects that altered splicing of *IKZF1* have on the expression of downstream target genes, but also, its potential association with the risk for autoimmunity.

## Material and methods

### Resource Availability

#### Lead Contact

Further information and requests for resources and reagents should be directed to and will be fulfilled by the lead contact, Dr Patrick Concannon.

#### Materials availability

All unique cell clones generated in this study are available from the lead contact with a completed material transfer agreement.

#### Data and code availability

RNA-seq and ATAC-seq data have been deposited in the National Center for Biotechnology Information Gene Expression Omnibus (NCBI GEO) under accession numbers GSE252707 (RNA-seq) and GSE255337 (ATAC-seq) and are publicly available as of the date of publication. Accession numbers are listed in the key resources table. Original western blot images have been deposited in the Figshare repository (**Supplemental Note**). All original code is available on GitHub (https://github.com/jrbnewman/IKZF1). Any additional information required to reanalyze the data reported in this paper is available from the lead contact upon request.

### Experimental model and subject details

#### Gene editing and clone selection

CRISPR-Cas9 technology was used to generate cloned lines from the parental Jurkat E6-1 cell line (ATCC TIB-152) with truncating mutations in exons 4 and 6 of the main *IKZF1* transcript (Ensembl ID ENST00000331340.8, Ensembl release 99). Two different sgRNAs were designed to target each exon at sites with limited homology to other IKAROS family members to avoid potential off-target effects. The sgRNAs, together with purified Cas9 protein (Dharmacon), in a ratio of 2.5:1 were individually electroporated into Jurkat E6-1 cells using the X-005 of the Nucleofector II (Lonza). Exon 4 and exon 6 were individually targeted in Jurkat E6-1 cells using Edit-R Modified Synthetic sgRNA (purified/deprotected/desalted) sequences sg1=GCGTGCCTGTGAAATGAATG or sg2=TTACGAATGCTTGATGCCTC (for exon 4), and sg1=CCACAACTACTTGGAAAGCA or sg2=CTTGGAAAGCATGGGCCTTC (for exon 6), (Dharmacon; Nuclease options: *streptococcus pyogenes* Cas9, PAM: NGG, Relative to target: after, Target length: 20, Cut relative to target 5’ start: Sense: 17 Antisense: 17, Repeat Sequence: GUUUUAGAGCUAUGCUGUUUUG, Relative to guide: 3’). Transfection efficiency was assessed by a T7 assay three days post-transfection (NEB EnGen® Mutation Detection Kit, Cat# E3321S). Cloned Jurkat cell lines were isolated by serial dilution in U-bottom 96-well plates and cultured for three weeks at 37°C (humidified atmosphere of 5% CO2) in 25mM HEPES RPMI 1640 (ThermoFisher, Cat#22400089) supplemented with 10% Fetal Bovine Serum, Sodium Pyruvate, GlutaMax and Penicillin/Streptomycin. Mutations in individual clones were characterized by PCR amplification and Sanger DNA sequencing on both strands. Selected clones, containing either mono-allelic or bi-allelic truncating mutations, were expanded in supplemented RPMI. For rigor, genomic DNA sequences were further confirmed in an additional separate DNA extraction performed on the same day as RNA and protein extractions. Additional clones containing truncating mutations in both exons were generated after targeting exon 4 with sg1 in one of the clones containing exon 6 truncating mutations. DNA sequences from clones with biallelic mutations were determined by cloning the specific blunt-ended PCR products in One Shot™ TOP10 Chemically Competent E. coli cells using Zero Blunt™ TOPO™ PCR Cloning Kit (ThermoFisher, Cat#K280020) following the manufacturer’s protocol. Heat-shock transformed bacteria were grown on LB supplemented agar plates containing 50 µg/ml kanamycin, selected colonies were expanded for 24 hours at 37°C with agitation, plasmid DNA was extracted using QIAprep Spin Miniprep kit (Qiagen, Cat#27106) and the SP6 promoter/primer site was used to prime DNA Sanger sequencing.

### Laboratory method details

#### RNA extraction and library preparation

RNA was isolated from selected clones (**S1 Table**) using the RNeasy Plus Mini kit (Qiagen Cat# 74134). RNA yield was measured by fluorometric quantitation using Qubit RNA Broad Range kit (ThermoFisher Cat# Q10210) and RNA integrity and purity were determined by automated electrophoresis (Agilent TapeStation system). RNA-seq libraries were prepared using NEBNext® Ultra II Directional RNA Library Prep Kit for Illumina (New England Biolabs Cat# E7760) according to manufacturer protocol.

#### Nuclei and ATAC-seq library preparation

Nuclei isolation, transposase incubation, and library preparation of *IKZF1*-targeted Jurkat clones was carried out following the Omni-ATAC-seq protocol (35). A total of 50,000 cells per clone were harvested and lysed in Base Buffer (10mM Tris-HCl, 10mM NaCl, 3mM MgCl2, pH 7.4) with 0.1% NP-40 (Sigma/Roche Cat# 11332473001), 0.1% Tween-20 (Sigma-Aldrich Cat# 11332465001) and 0.01% Digitonin (Promega Cat# G9441). After pelleting the cells, a transposition reaction was performed using Nextera DNA Library Prep Kit (Illumina, Cat# FC-121-1030). PCR amplification of transposed DNA fragments was performed incorporating primers designed based on Omni-ATAC-seq protocol (36)and NEBNext High-Fidelity 2x PCR Master Mix (New England Biolabs Cat# M0541L). Library purification was achieved by using Agencourt AMPure XP beads (Beckman Coulter, Cat# A63881). Library yield was determined using the Qubit High Sensitivity dsDNA (ThermoFisher, Cat# Q32851), and DNA integrity was assessed by D1000 High Sensitivity Tape Station (Agilent TapeStation system). Libraries were sequenced on a NextSeq 500 (Illumina) as a 75bp paired-end sequencing run.

#### Immunoblotting

Protein was extracted from whole cell lysates using extraction buffer containing 0.05 M Tris HCl pH 8.0, 0.12 M NaCl, 0.5% NP40, 1 mM EDTA, 0.05 M NaF, 1mM Na3VO4, 1 mM β-mercaptoethanol, cOmplete mini-protease inhibitor cocktail and PhosSTOP phosphatase inhibitor tablets (Roche Cat# 4693159001 and Cat# 4906837001). Protein (25 µg) was denatured with LDS Sample Buffer and Sample Reducing Agent and separated on 4-12% Bis-Tris 1 mm mini protein gels using MES SDS Running Buffer with 0.1% NuPAGE Antioxidant (Invitrogen Cat#NP0005). Molecular weight was assessed using the eBlue Plus2 Prestained Protein Standard (Invitrogen). Protein was transferred to a polyvinylidene difluoride membrane (IPVH20200 Immobilon) on a Novex XCell II Blot Module using Transfer Buffer (Invitrogen) containing 10% Methanol and 0.1% Antioxidant. Membranes were blocked using 5% skim milk powder in Tris-Buffered Saline with 0.05% Tween-20 and incubated at a concentration of 1:2000 with rabbit recombinant anti-IKAROS (clone EPR13791) C-terminal antibody (Abcam Cat# ab190691), which recognizes seven isoforms with the predicted molecular weights of 58kDa, 48kDa, 48kDa, 43kDa, 41kDa, 32kDa and 53kDa; or rabbit polyclonal anti-IKAROS N-terminal antibody (Abcam Cat# ab229275), which targets amino acids 1-165 of the human isoform protein. Additional immunoblotting assays were performed using rabbit polyclonal anti-ZNFN1A2/HELIOS antibody at a concentration of 1:2000 (Abcam Cat# ab129434), rabbit monoclonal recombinant anti-Integrin beta 1 (EPR1040Y) antibody at a concentration of 1:1000 (Abcam Cat# ab134179), mouse polyclonal anti-IKZF3 antibody at a concentration of 1:1000 (Abcam Cat# ab88513) or rabbit monoclonal recombinant anti-IL-15 [EPR22635-214] antibody at a concentration of 1:1000 (Abcam Cat# ab273625). A mouse monoclonal anti-γ-tubulin (clone GTU-88) antibody (1:5000, Sigma-Aldrich Cat# T5326) was used to probe all blots as a protein loading control. Horse Radish Peroxide-conjugated goat anti-rabbit and anti-mouse IgG secondary antibodies (1:5000, BD Bioscience) were used for protein detection. Protein expression was analyzed by chemiluminescence using a GE Life Sciences AI600 imager.

### Quantification and statistical analysis

#### RNA sequencing and read processing

RNAseqlibraries were prepared according to Illumina protocols and sequenced with 2 × 151 nt paired reads on an Illumina HiSeq 3000 instrument (2.45 ×10^8^ ± 3.52 ×10^7^ paired reads/sample). Quality of sequencing data was assessed using GC content, and the percentage of adapter content, duplication rate, and homopolymer content in each sample (https://www.bioinformatics.babraham.ac.uk/projects/fastqc/). Illumina TruSeq adapter sequences were trimmed from read pairs using CutAdapt (version 2.8). Read pairs whose sequences overlapped by 37 nt or more (i.e. 25% of the read length) were merged into a single read using the BBMerge function of the BBMap tool suite (version 38.44; https://sourceforge.net/projects/bbmap/). Merged and unmerged adapter-trimmed reads that were less than 75 nt in length were then removed. Duplicate reads/read pairs were then removed from the length-filtered reads. The resulting set of processed sequence reads were used for quantification.

#### Quantification of gene expression

We employed a two-pass alignment strategy for estimating gene and transcript expression strategy to obtain the best probable set of expressed transcripts in Jurkat cells. For the first pass, we utilized Event Analysis (37) to annotate genes and transcripts and the human transcriptome (Ensembl release 99) in terms of exons, exonic sequence fragments based on transcript membership, and exon-exon junctions. Junction sequences were quantified by aligning reads to a catalog of all plausible junction sequences using Bowtie (version 1.2.2, (38)). Exonic sequences were quantified by aligning reads to the human genome version GRCh38.p13 using BWA-MEM (version 0.7.15, https://arxiv.org/abs/1303.3997). Genomic-mapping read alignments were intersected with a BED file of the genomic regions demarcating exonic sequences to calculate read coverage. Coverage was calculated as the average depth per nucleotide (APN = sum of read depth / length of feature). We defined exonic and junction sequences as detected if there was at least 1 mapped read in 50% of clones. We included transcripts from the human reference transcriptome for the first-pass quantification only if all their constituent features (exonic sequences and junctions) had at least 1 mapped read in 50% of clones. We then utilized RSEM (version 1.2.28) to quantify the reduced reference transcriptome in all clones presented in this study (39). Transcriptome references were prepared with ‘*rsem-prepare-reference*’ using the set of transcript sequences as FASTA sequence and a tab-delimited gene-to-transcript file as input (39). Default settings were used for all parameters. Transcript and gene abundance was estimated as transcripts per million (TPM) (40). A transcript was considered expressed if the TPM > 0 for >50% of all clones. We considered a transcript detected if it had a TPM>0 in at least 50% of all clones, and classified a gene as "expressed" if at least one of its annotated transcripts were detected.

For the second-pass quantification, we removed from our transcriptome reference annotation any transcript considered not detected in the first-pass quantification and any gene that did not have any detected transcripts. We then re-quantified this further reduced reference annotation using RSEM (same parameters as above). As with the first-pass quantification, a transcript was considered detected if it had a TPM > 0 in at least 50% of all clones, and a gene was considered "expressed" if at least one of its annotated transcripts were detected. Unless otherwise indicated, only gene-level estimates were used in subsequent analyses. Gene expression estimates were upper-quartile normalized and log-transformed due to overall performance of this strategy (23, 41, 42).

#### ATAC-seq read processing and peak calling

ATAC-seq libraries were sequenced using 76-bp paired-end reads on an Illumina NextSeq 550. Sequencing reads were processed using the PEPATAC pipeline (43). Briefly, reads were trimmed using Skewer (version 0.2.2) (44), and reads that mapped to mitochondrial or human repeat regions were removed. The remaining reads were mapped to the complete human genome version GRCh38.p13 using BWA-MEM (https://arxiv.org/abs/1303.3997). PCR duplicates were removed using SAMBLASTER (45). Enzymatic cut sites were inferred from read alignments and peaks were called using MACS2 (https://www.biorxiv.org/content/10.1101/496521v1). Data quality was assessed using TSS enrichment plots, and plots of the fraction and cumulative fraction of reads in genomic features to evaluate enrichment of reads in genomic features (43).

Consensus peaks were determined by identifying overlapping (1 bp) peaks between all clones and the overlapping peak with the highest score was used as the consensus peak coordinate. Peaks present in at least 2 clones with a score of ≥ 1 per million were retained. Consensus peak counts for each clone were calculated by multiplying individual peak counts for an overlapping peak by the percent overlap of the sample peak with the consensus peak (43).

Peaks with fewer than 10 reads in ≥ 50% of samples were excluded from further analysis. Peak count normalization was performed using the trimmed mean of M-values (TMM) method (46) implemented in the edgeR R package (47). Linear modeling of peak counts was enabled by performing a mean-variance modeling-based transformation using limma-voom (48). Outlier peaks were identified by clustering sample counts for each peak (one at a time using *k*-means clustering, with *k* = 2 clusters); any peak in an individual sample that clustered separately from all other samples was considered an outlier peak and excluded from further analysis. Consensus peaks were annotated using HOMER annotatePeaks.pl script (49).

#### Differential expression and accessibility analysis

To assess differential gene expression, for each gene, gene expression estimates were modeled as *Y_ij_* ∼ *µ* + *g_i_* + *ε_ij_*, where *Y_ij_* is the log-transformed normalized transcripts per million of group *i*, clone *j*; *g_i_* is group; and *ε_ij_* is the residual, distributed ∼N(0,σ^2^). We utilized a similar model to test differential accessibility, where for each peak, the peak count data were modeled as *Y_ij_*∼ *µ* + *g_i_* + *ε_ij_*, where *Y_ij_*is the log-transformed normalized counts per million of group *i*, clone *j*; *g_i_* is group; and *ε_ij_* is the residual, distributed ∼N(0,σ^2^). In terms of group membership of clones, we devised several arrangements of samples to test different hypotheses. The first hypothesis tested was that editing of exons 4 and/or 6 of *IKZF1* will result in changes in changes in transcription and chromatin remodeling in Jurkat cells. This was a binary “WT” and “targeted” comparison, where clones were organized into “WT” and “targeted”. The second hypothesis tested was that the editing of different exons (ie. exon 4, exon 6, or exons 4 and 6) results in different changes in transcription and chromatin remodeling in Jurkat cells. Clones were grouped by targeted exon(s) (“WT”, “exon 4”, “exon 6”, “exon 4 and 6”). The third hypothesis tested was that the detectable level of IKAROS protein influences changes in gene expression and chromatin accessibility. Clones were grouped by protein detection on a Western blot and by targeted exon(s): “WT”, “exon 4 low-expressing”, “exon 4 high-expression”, “exon 6”, “exon 4 and 6”. The fourth hypothesis tested was that the number of N-terminal zinc finger domains that can be potentially encoded in functional IKAROS isoforms, by way of the number of exons targeted, determines gene expression and chromatin accessibility. Clones for this test were grouped into “WT”, “single-exon targeted”, and “double-exon targeted”. Details of group membership for each clone can be found in **S2 Table**. For each hypothesis, the mean gene expression or mean peak height was determined for each group, and the log_2_-transformed fold change calculated for each pairwise comparison of groups of clones.

#### Gene set enrichment analysis

Enrichment tests targeting sets of autoimmune disease risk genes were performed using a two-sided χ^2^ test. Gene sets from three recent T1D GWAS (9, 50, 51) were compiled to test enrichment for T1D risk genes (N = 231 genes total). Risk genes for SLE were derived from several sources (52, 53) and combined to test enrichment for SLE risk genes (N = 681 genes total). To test for enrichment of autoimmune disorder risk genes broadly, we combined gene lists from catalogs of risk loci for multiple autoimmune disease (52, 53) (N = 2,901 genes). For all tests, binary response variables were used to indicate gene membership (0=not in gene set; 1=in gene set) and statistical significance of a gene (0=not significant; 1=significantly differentially expression and/or has a proximal differentially accessible peak).

## Acknowledgements

The authors thank Jason Smith and Nathan Sheffield for their assistance with the PEPATAC pipeline and analyses of ATAC-seq data.

## Supportive Information

### Supplemental Note

**S1 Table. Characteristics of Jurkat clones derived from CRISPR editing of *IKZF1***

**S2 Table. Group membership of Jurkat clones**

**S1 Fig. Characteristics of *IKZF1* isoforms and the impact of gene-editing events.** Indicated are the lengths in amino acids (aa), molecular weights (kDa), database IDs and structures of documented isoforms of *IKZF1*/Ikaros that include both the DNA binding and dominant-negative isoforms. Potential expression indicates which Ikaros isoforms can be produced in Jurkat clones with editing events in exon 4, exon 6 or in both. WT = wild-type; EX4 = exon 4; EX6 = exon 6; EX6+4 = both exons. Exons alternatively spliced among the reported IKAROS functional isoforms are colored orange. Exons containing zinc fingers but that are not alternatively spliced are indicated in red. All other exons are colored blue.

**S2 Fig. IKAROS protein expression in targeted clones.** IKAROS protein expression as assayed by immunoblotting of clones with targeting events in exon 4 (A), exon 6 (B) and both exons (C), as compared to wild type clones (WT), using a primary antibody specific for the C-terminal end of the protein. The same blots were stripped and probed with an antibody against γ-tubulin as protein loading control. Black arrows indicate the expected sizes of IKAROS isoforms containing the N-terminal zinc fingers encoded by exon 5 (i.e. Ik-1, Ik-2, Ik-2a, Ik-3, Ik-3a, Ik-4; S Figure 1).

**S3 Fig. Differentially accessible ATAC-seq peaks in intergenic regions.** (A) Intergenic ATAC-seq peak analyzed in each comparison made. (B) Overlap of all intergenic DAPs between WT, single-targeted, and double-targeted clones.

**S4 Fig. Upset plot of comparisons between DE and DA genes between the three groups of clones.** Gene membership in each category (DA genes: WT vs single-targeted, WT vs double-targeted, single-targeted vs double-targeted; DE genes: WT vs single-targeted, WT vs double-targeted, single-targeted vs double-targeted) is indicated by black dots. If genes belong to multiple categories (e.g. DA WT vs single-targeted and DE WT vs single-targeted) vertical black lines are drawn connecting each category. The total number of genes in each of the six categories is presented by the horizontal bar chart to the left of category labels.

**S5 Fig. Effects of *IKZF1* editing on example genes, *IKZF2* (A), *IKZF3* (B), *ITGB1* (C), and *IL15* (D), in terms of chromatin accessibility (i), gene expression (ii), and protein levels (iii).** All changes in chromatin accessibility associated with each gene are presented here. Red boxplots = WT clones; white boxplots = single-exon clones; purple boxplots = double-exon clones; pink/red dots = WT clones; green dots = low protein-expressing exon 4-targeted clones; yellow dots = high protein-expressing exon 4-targeted clones; blue dots = exon 6-targeted clones; purple dots = double-exon clones.

